# HUHgle: An Interactive Substrate Design Tool for Covalent Protein-ssDNA Labeling Using HUH-tags

**DOI:** 10.1101/2024.03.15.585203

**Authors:** Adam T. Smiley, Natalia S. Babilonia-Díaz, Aspen J. Hughes, Andrew C.D. Lemmex, Michael J.M. Anderson, Kassidy J. Tompkins, Wendy R. Gordon

## Abstract

HUH-tags have emerged as versatile fusion partners that mediate sequence specific protein-ssDNA bioconjugation through a simple and efficient reaction. Here we present HUHgle, a python-based interactive tool for the visualization, design, and optimization of substrates for HUH-tag mediated covalent labeling of proteins of interest with ssDNA substrates of interest. HUHgle streamlines design processes by integrating an intuitive plotting interface with a search function capable of predicting and displaying protein-ssDNA bioconjugate formation efficiency and specificity in proposed HUH-tag/ssDNA sequence combinations. Validation demonstrates that HUHgle accurately predicts product formation of HUH-tag mediated bioconjugation for single- and orthogonal-labeling reactions. In order to maximize the accessibility and utility of HUHgle, we have implemented it as a user-friendly Google Colab notebook which facilitates broad use of this tool, regardless of coding expertise.

**Graphical Abstract:** 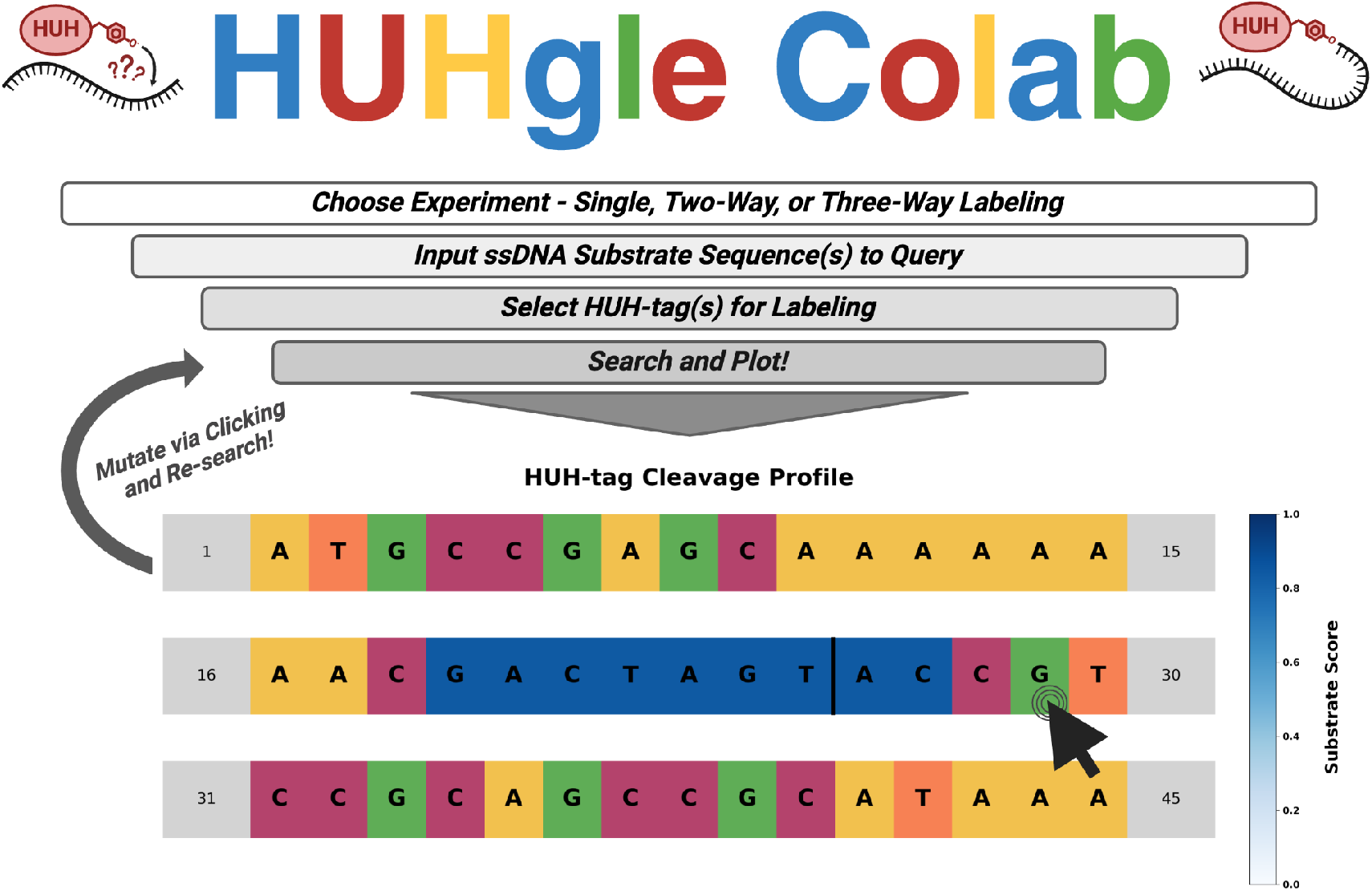

## Introduction

HUH-endonucleases are a diverse group of mechanistically similar enzymes that contain a unifying metal-coordinating motif most often comprised of a pair of histidines (H) separated by a hydrophobic residue (U) [1,2]. The HUH superfamily includes nucleases that mediate a host of biological processes, ranging from replication, to conjugation, and even transposition [1-7]. These enzymes mediate their functions through a common single-stranded DNA (ssDNA) processing mechanism in which a coordinated divalent cation polarizes the phosphate backbone of the substrate to enable the catalytic tyrosine to cleave and covalently bind to the substrate’s newly exposed 5’ end via a phosphotyrosine linkage (Figure 1A) [1]. Though these covalent protein-ssDNA adducts are transient products in endogenous contexts, bioconjugates formed by reacting recombinant HUH-endonucleases with ssDNA substrates harboring their specific cleavage motifs are long-lived and non-labile [2,8].

**Figure 1.**
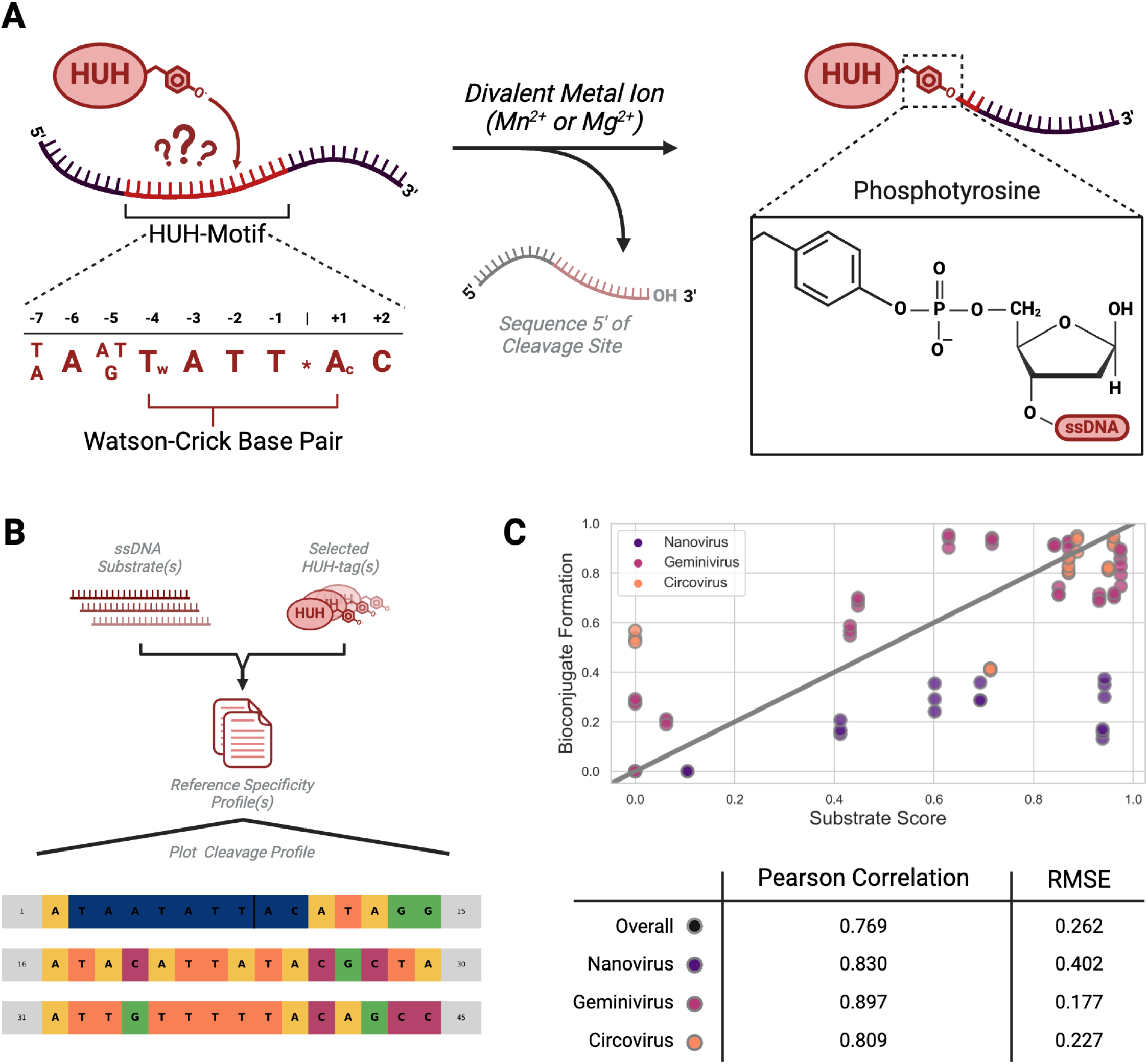
Introduction to HUH-tags and HUHgle: (A) Graphical schematic of an *in vitro* HUH-tag bioconjugation reaction. (B) High-level graphical overview of the HUHgle substrate design tool. (C) Scatterplot depicting the relationship between substrate score and bioconjugate formation calculated via quantification of *in vitro* HUH-tag reactions in triplicate. Data are colored by viral family with associated Pearson correlation and root mean squared error statistics at the bottom. Individual reaction gels that make up the data from (C) can be found in figure S2 in the supporting information.

‘HUH-tags’ have emerged as a versatile bioconjugation platform to mediate simple, efficient, and sequence-specific linkage of proteins to ssDNA substrates (Figure 1A) [8]. Similar to the SNAP-, CLIP-, and HALO-tag systems, HUH-tags are self-labeling enzymes that covalently link to their substrates [9-11]. HUH-tags are a superb option for protein-DNA bioconjugation because, in contrast to other self-modifying enzymes, they react directly with ssDNA oligonucleotides harboring an appended 5’ HUH-tag cleavage site (∼9 bases) without the need for expensive chemical modifications. HUH-tags have broad applicability in a diversity of biotechnology applications. For example, these enzymes have found use in novel genome engineering systems, particularly CRISPR-based tools. They have been used to tether DNA templates to Cas9 in order to improve homology directed repair efficiency and gene knock in [12-14]. Additionally, HUH-tags play a critical role in ‘click editing’ [15], an emerging alternative to prime editing [16], in which an HUH-endonuclease covalently tethers a ssDNA template encoding user-specified mutations to a Cas9-Klenow fragment fusion for site-specific rewriting of genetic information. Beyond genome engineering applications, HUH-tags have been utilized in a user-friendly molecular tension sensing tool called RAD-TGTs (“Rupture And Deliver Tension Gauge Tethers”) [17], which provide a simple platform for characterizing cellular tension profiles. Moreover, HUH-tags have enormous potential as a facile means by which to covalently link user-specified DNA sequences to binding proteins for sequencing-based epitope profiling via DNA barcoding [18].

The two classes of HUH-endonucleases deployed as HUH-tags are the replication initiator proteins (Reps), which are involved in rolling circle replication in ssDNA viruses and plasmids, and the relaxases, which are involved in plasmid conjugation in prokaryotes [1-4, 8,9]. While both classes have been explored as HUH-tags, the Reps – particularly those from phylum *Cressdnaviricota* [19] – have garnered particular popularity as fusion partners. This is owing to their compact size, their robust reactivity, and their concise cleavage motifs [2]. Indeed, Rep HUH-tags derived from this phylum exhibit a somewhat common specificity – cleavage sites follow roughly a ‘TAATATT^*^AC’ nonanucleotide motif, where cleavage occurs at the ‘^*^’ and the covalent bond is formed to the +1 adenosine (Figure 1A) – but there is variability in specificity profile among the families in this phylum [2]. This variability in specificity has proven to be both an advantage and disadvantage: it enables the identification of intrinsic orthogonality for simultaneous, non-cross reactive labeling of multiple HUH-tags to their respective ssDNA substrates in a single reaction, but it also introduces complexity into the substrate design process, making the prediction of low-efficiency ‘off-target’ cleavage sites difficult without computational tools [2]. To address these challenges and to enhance the accessibility of HUH-tags, we have developed HUHgle (Figure 1B), a computational tool specifically designed to streamline substrate design and make HUH-tag applications more intuitive and approachable to all.

## Results & Discussion

### HUHgle - The HUH-tag Substrate Design Tool

In a previous study, we characterized the specificity of a panel of HUH-tags using a deep-sequencing method, ‘HUH-seq’ [2]. We showed this assay produces sequence logos indicative of an HUH-tag’s specificity profile that were in agreement with cognate origin of replication sites (*ori*) (Figure S1) [2]. Though the assayed HUH-tags cleave similar *ori* sequences endogenously, our profiling revealed a diversity of promiscuous activity across these enzymes. Though not the initial intention of this assay, we speculated that this data could be used to predict bioconjugate formation efficiency for a given HUH-tag/ssDNA substrate pair. Thus, we developed a ‘scoring’ metric from this data by which a substrate is scored between 0 - 1 based on the extent to which it can be acted upon by a given HUH-tag. We performed *in vitro* bioconjugation reactions with our panel of eight HUH-tags on substrates with a diversity of scores (Figure 1C, S2). We found that substrate score has a high positive correlation with bioconjugate formation efficiency across all assayed HUH-tags, with an overall pearson correlation coefficient of 0.769 (Figure 1C). When grouping the HUH-tags into their families within phylum *Cressdnaviricota* – specifically, *nano-, gemini-*, and *circoviridae* – correlation coefficients improve to 0.830, 0.897, and 0.809, respectively, demonstrating high functional similarity within a family. Additionally, we found that in some families substrate score can even be predictive of total bioconjugate yield. Specifically, we calculated the root-mean-square error between this metric and percent bioconjugate formation to be as low as 0.177 in HUH-tags from *geminiviridae* (Figure 1C). Overall, our substrate score metric is a strong predictor of a sequence’s performance in an HUH-tag/ssDNA substrate pair and serves as the foundation of the HUHgle substrate design tool.

Briefly, HUHgle is a python-scripted search function followed by interactive plotting, developed to simplify the process of substrate design and optimization for HUH-tag applications. Users begin by specifying their experimental approach – either single-, two-, or three-way bioconjugation. Upon inputting desired ssDNA substrate sequence(s), the user is prompted to select their HUH-tag(s). HUHgle has a panel of eight HUH-tags to choose from (supplementary table 1), each with their own unique specificity profile. The user can select their own enzyme from the panel or HUHgle can recommend three HUH-tags based on minimizing predicted interaction with the substrate to limit off-target product formation. HUHgle can append a high efficiency site of interaction on the 5’ end of the input substrate based on the selected enzyme, or the user can design this site by hand. Following this, HUHgle’s search function evaluates the sequence against the specificity profile of the selected HUH-tag. Subsequently, it plots a cleavage profile for the substrate, specifically highlighting the efficiency and location of potential cleavage sites by recoloring them with a monochromatic gradient indicative of substrate score (Figure 1B). This plot has an interactive interface that allows for *ad hoc* modification of the substrate sequence by directly clicking on the bases to cycle through the different options – each alteration is automatically reassessed and plotted, enabling rapid iteration towards an optimized substrate sequence.

### HUHgle Accurately Identifies HUH-tag Mediated Protein-ssDNA Bioconjugate Formation

HUHgle plots demonstrate the differences in specificity across members of our panel of HUH-tags (Figure 2). Though variation in cleavage profiles of our HUH-tags are seemingly modest at first glance (Figure S1), these minor changes can yield dramatic differences in bioconjugation results on the same substrate. For example, in a simple single-labeling bioconjugation reaction with HUH-tags from either the geminivirus wheat dwarf virus (WDV), the circovirus porcine circovirus 2 (PCV2), or the nanovirus faba bean necrotic yellow virus (FBNYV), we show that HUHgle predicts dramatically different cleavage profiles on the same substrate across these three enzymes (Figure 2). The designed cleavage site (highlighted with a red box) is the highest scored site of interaction across all three selected HUH-tags, but the substrate ranges from ideal for WDV which is predicted to interact at a single site (Figure 2A), to suboptimal for FBNYV which is predicted to have three additional low-scored sites (Figure 2C), and even poor for PCV2 which is predicted to have two additional modest-scored sites (Figure 2B). To validate HUHgle’s predictions, we performed *in vitro* bioconjugation reactions with recombinantly expressed HUH-tags and synthetic oligonucleotides. Because HUH-tags form covalent bonds to their substrates, bioconjugation reaction efficiency and specificity can be analyzed via SDS-PAGE through quantification of bands with increased molecular weight. Surprisingly, off-target product formation is only prominent in conditions with sub-stoichiometric amounts of substrate, even in seemingly unfavorable reactions with PCV2 (Figure 2). This suggests that high scoring cognate-like sites of interaction can mask low scoring sites through preferential or enhanced association. Engineering the designed cleavage site to have a lower score dramatically increases the amount of off-target interaction in an otherwise identical substrate across these three HUH-tags, regardless of the reaction stoichiometry (Figure S3). Overall, our data indicates that HUH-tag choice can be non-trivial and HUHgle is capable of analyzing a sequence of interest and recommending enzymes that minimize off-target product formation.

**Figure 2.**
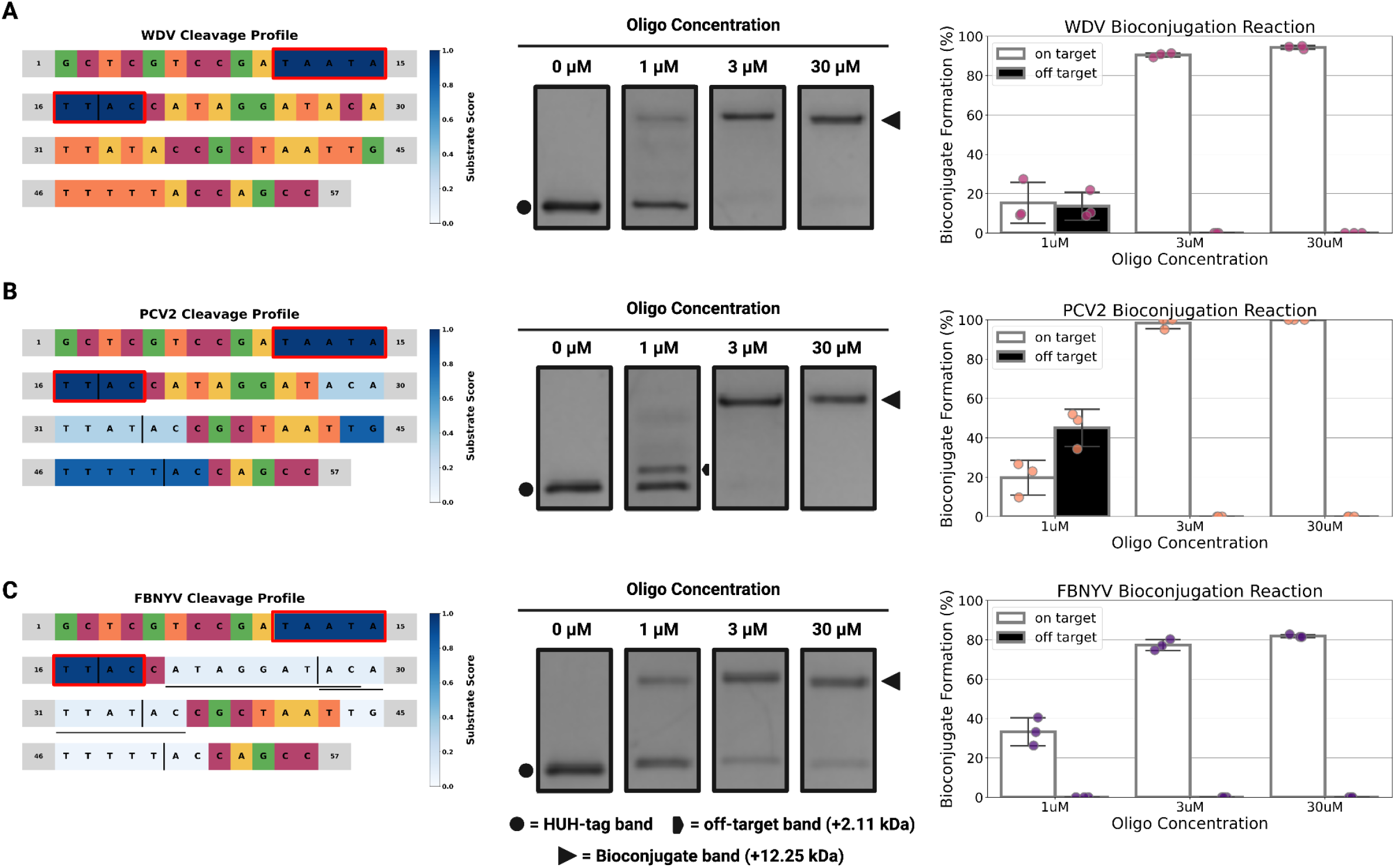
HUHgle predictions in one-way bioconjugation: Subfigures (A), (B), and (C) show HUHgle substrate interaction plots (left), *in vitro* HUH-tag bioconjugation reactions visualized via SDS-PAGE (center), and barplots quantifying these reactions (right) for the HUH-tags WDV (A), PCV2 (B), and FBNYV (C) on the substrate *HUHgle5* (see supporting information). Reactions were performed in final concentrations of 3 μM HUH-tag and indicated concentration of oligo in 50 mM HEPES pH 8.0, 50 mM NaCl, 1 mM DTT, and 1 mM MnCl_2_ for one hour at 37°C. HUH-tag bands, on-target bioconjugation bands, and the most prominent off-target bioconjugation are indicated with unique black characters – see the legend at the bottom of the figure.

### HUHgle is an Effective Tool in the Design of Orthogonal Bioconjugation Reactions

There is sufficient variation in sequence specificity profiles across our panel of HUH-tags to enable exclusive, non-cross reactive labeling of pairs, and even triplets, of enzymes. We used our substrate scoring metric to identify and rate combinations of HUH-tags based on their number of mutually exclusive high-scoring sequences. Using this information, HUHgle can aid in the design of two- or three-way orthogonal labeling reactions by recommending optimal HUH-tag pairs/triplets for a given set of substrates and appending non-cross reactive sites of interaction to their 5’ ends to enable orthogonal bioconjugation. To demonstrate this, we used our tool to design a series of two- and three-way labeling reactions. Briefly, we used HUHgle to design a simple orthogonal bioconjugation reaction in which two substrates are exclusively acted upon by their respective HUH-tag (Figure 3A). We input arbitrary ssDNA sequences into HUHgle and our tool recommended a number of combinations capable of mediating our desired orthogonal labeling from which we chose a pair from duck circovirus (DCV) and tomato golden mosaic virus (TGMV), members of *circo-* and *geminivirdae*, respectively. These enzymes compose a strong orthogonal pair with a diversity of mutually exclusive interaction sites across a range of scores (Figure 3B). HUHgle appended the highest scoring orthogonal sequences from this pair to the 5’ ends of each of our substrates and plotted their predicted reaction profiles (Figure 3A). Subsequent *in vitro* reactions validates these predictions by demonstrating high substrate selectivity and bioconjugate formation efficiency in reactions with one-to-one substrate-to-enzyme stoichiometry (Figure 3A & C). We expand on two-way labeling by designing a single substrate harboring orthogonal interaction sites for enzymes from banana bunchy top virus (BBTV) and cabbage leaf curl virus (CLCV), members of *nano-* and *geminivirdae*, respectively (Figure 3D, E & F). Subsequent *in vitro* reactions with this single substrate and HUH-tag pair validates our tool’s plot, again demonstrating high selectivity and bioconjugate formation efficiency (Figure 3D & G). Furthermore, to demonstrate HUHgle’s capabilities in three-way labeling, we used our tool to design substrates to be exclusively acted upon by their respective HUH-tag (Figure 4A). Upon entering our set of sequences, HUHgle analyzed them and recommended a list of orthogonal HUH-tag triplets, from which we selected the set of BBTV, DCV, and WDV. These enzymes compose a strong orthogonal triplet with a diversity of sequences (Figure 4C & B), from which HUHgle identified the highest scoring orthogonal sites and appended them to the 5’ end of our sequences. *In vitro* reactions again validate the substrate selectivity designed by HUHgle (Figure 4B). Finally, we expand on three-way labeling by designing a single substrate which harbors three mutually exclusive interaction sites for an additional orthogonal triplet of HUH-tags, BBTV, DCV, and TGMV, and again validate HUHgle’s predictions via *in vitro* reaction and analysis (Figure S4).

**Figure 3.**
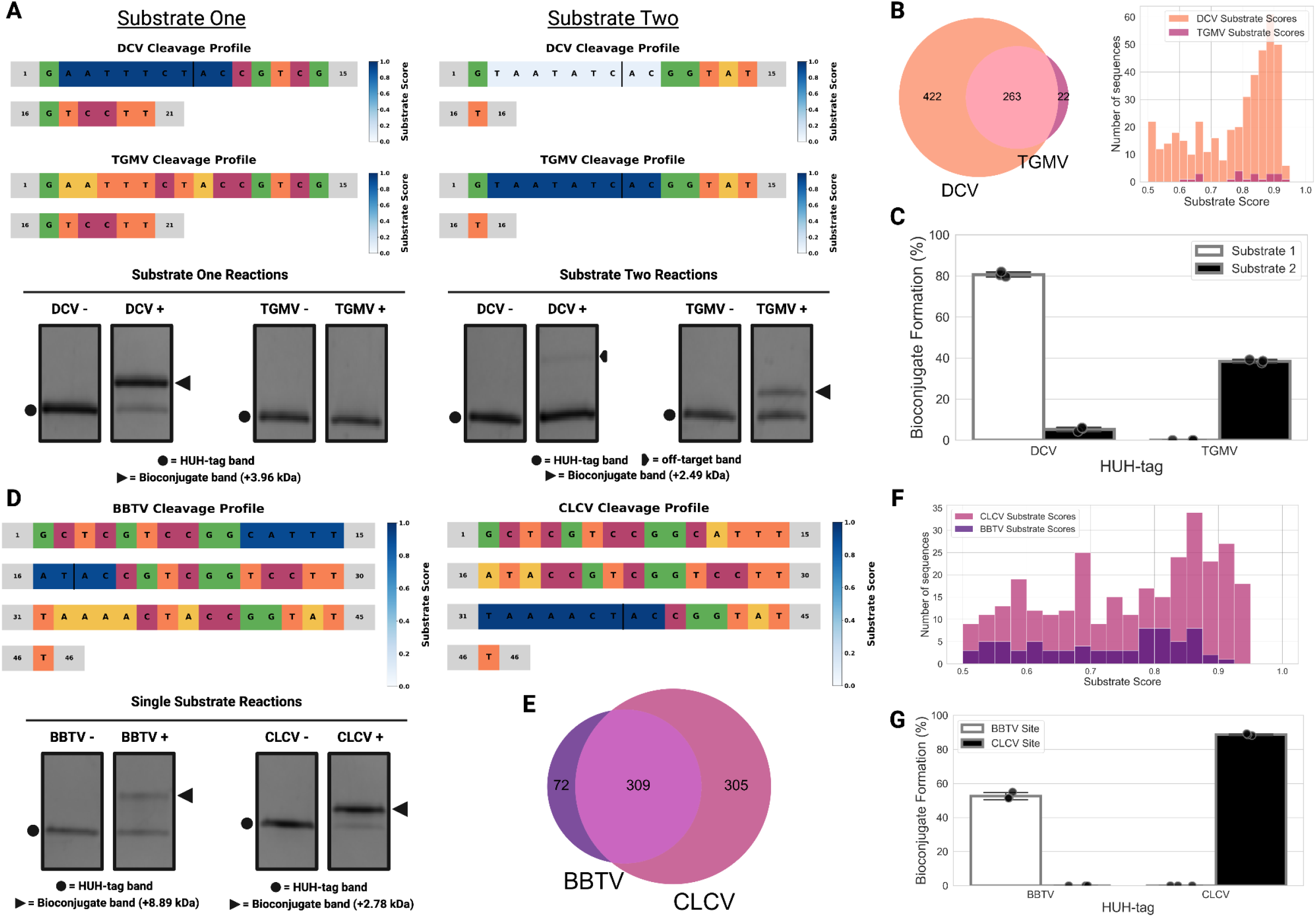
HUHgle predictions in two-way orthogonal bioconjugation. Subfigures (A) and (D) show HUHgle substrate interaction plots indicating cleavage on a designed orthogonal substrate pair (A) and single oligo containing two interaction sites (B) (top), and *in vitro* HUH-tag bioconjugation reactions with these substrates visualized via SDS-PAGE (bottom) for the HUH-tag pairs DCV and TGMV, and BBTV and CLCV, respectively. Subfigures (C) and (G) show barplots quantifying these reactions for the labeled HUH-tag pairs. Subfigures (B) and (E) show Venn diagrams indicating the number of orthogonal and non-orthogonal substrates scoring above 0.5 across the indicated pair. The histograms in subfigures (B) and (F) indicate the number of exclusively orthogonal substrates binned by substrate score across the indicated pair.

**Figure 4.**
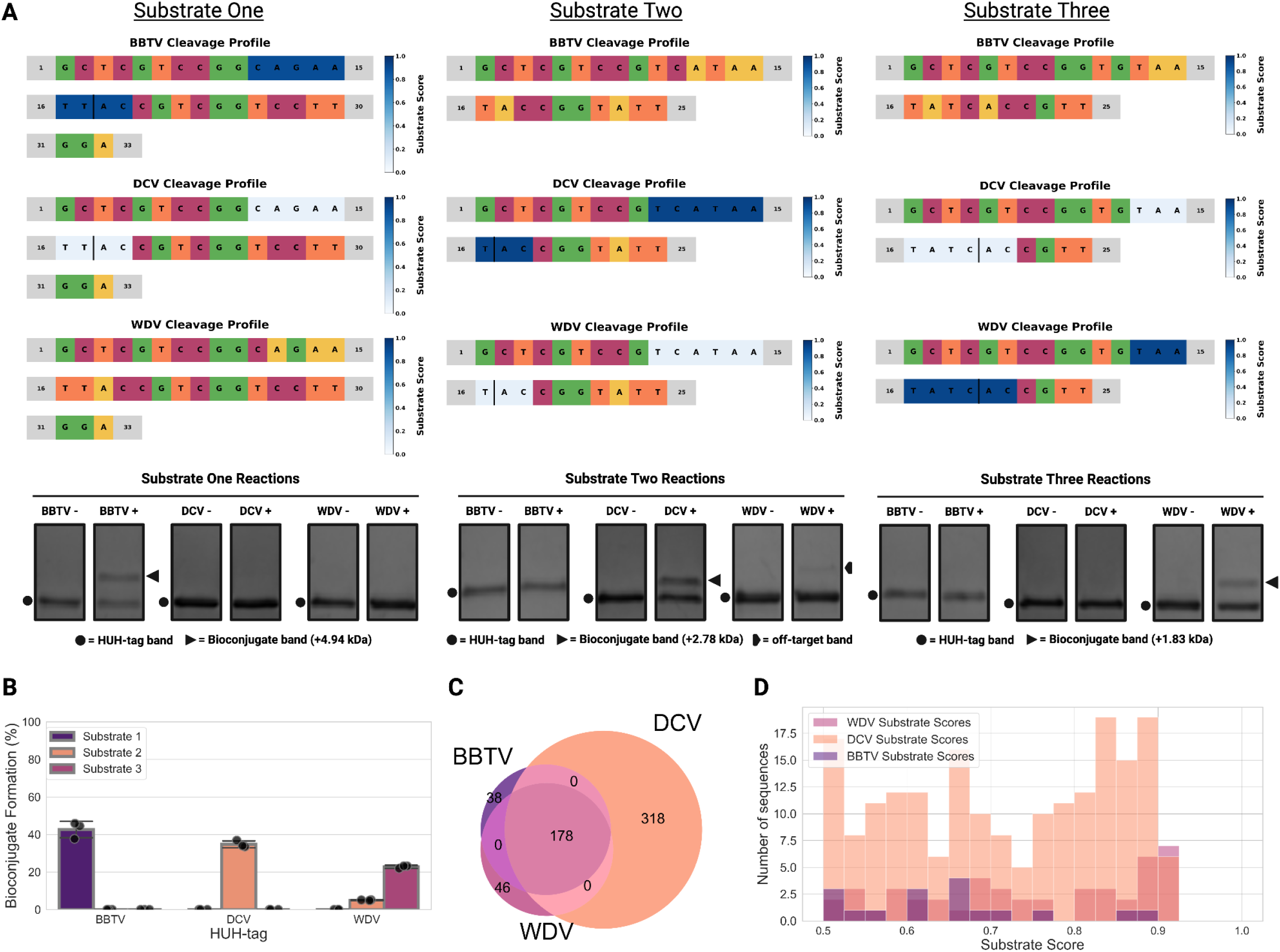
HUHgle predictions in three-way orthogonal bioconjugation. Subfigure (A) shows HUHgle substrate interaction plots indicating cleavage on a designed set of three orthogonal substrates (top), and *in vitro* HUH-tag bioconjugation reactions with these substrates visualized via SDS-PAGE (bottom) for the orthogonal combination of the HUH-tags BBTV, DCV, and WDV. Subfigure (B) shows a barplot quantifying these reactions for the labeled HUH-tag and substrate from the orthogonal set. Subfigure (C) shows a Venn diagram indicating the number of orthogonal and non-orthogonal substrates scoring above 0.5 across the indicated set of HUH-tags. Subfigure (D) shows a histogram that indicates the number of exclusively orthogonal substrates binned by substrate score across the indicated set of HUH-tags.

We note that two- and three-way reactions have modest yields in comparison to one-way reactions. This is likely due to slower kinetics on the sub-optimal substrates that enable orthogonality. However, this can be overcome by increasing reaction time or substrate concentration. For example, three-way reactions in Figure 4 were incubated for one hour and yielded 30-40% bioconjugate formation, whereas the comparable three-way reactions in Figure S4 were incubated for 24 hours and yielded ∼80% bioconjugate formation.

### HUHgle Bonus Features

HUHgle has three ‘bonus’ features that we find occasionally useful in the design of HUH-tag substrates. First, HUHgle can broaden its sequence search space through *in silico* prediction based on the recent discovery that HUH-endonucleases from phylum *Cressdnaviricota* have more plasticity in sequence specificity than initially identified. Specifically, these nucleases are dependent on Watson-Crick base pairing between positions -4 and +1 within their substrates regardless of the combination of nucleotides that form the pairing [20]. Our deep sequencing data was collected on substrates containing a constant ‘AC’ dinucleotide in the +1 and +2 positions, limiting our resolution on this intramolecular dependency until recently [2, 20]. We show the potential usefulness of this ‘expanded mode’ through comparative HUHgle plotting and subsequent *in vitro* reaction of WDV with an oligonucleotide containing a number of cryptic cleavage sites that are revealed by this feature (Figure 5A). While occasionally useful, we consider expanded mode to be a bonus feature because while this substrate expansion works as a general rule, there are some exceptions and some non-WC combinations can enable cleavage, depending on the specific HUH-tag. Second, HUHgle can predict interactions on substrates containing nucleotides corresponding to IUPAC ambiguity codes. This feature takes in the partially randomized substrate, generates all possible variants, and then analyzes them for potential sites of interaction. Following this, the identified sites are plotted as a histogram of interaction sequences binned by substrate score. Though unnecessary in most HUH-tag applications, we see value for this bonus feature in the context of DNA barcoding [18]. Indeed, barcoding strategies often employ a randomized ‘Universal Molecular Identifier’ (UMI) to mediate the removal of PCR duplicates to increase the accuracy of epitope/variant quantification. We used this tool to design a barcoding substrate with minimal off-target interaction for HUH-tag mediated barcoding applications. Conventional randomized UMI sequences are predicted to have thousands of potential interaction sites whereas UMI sequences with intervening ‘G’ bases almost completely eliminate these putative off-target interactions (Figure 5B). Third, we designed a bonus feature to identify potential sites of interaction in sequences that are too large to be visualized as standard HUHgle plots. This feature enables HUHgle to exhaustively search a .FASTA file for potential sites of interaction and plot them as a histogram of sequences binned by substrate score. This bonus feature is able to make predictions on both linear and circular sequences. One potential use case of this feature is in the prediction of cognate *ori* sites in genomes belonging to phylum *Cressdnaviricota*. Because these sites form stem loop structures to increase HUH-endonuclease accessibility, we implemented an optional folding step using ViennaRNA to better classify identified cleavage sites as *ori* sequences [21]. As proof of concept, we used our tool to identify sites of interaction, and subsequently predict and fold the likely *ori* sequence, from the genome of a member of our panel of HUH-tags, FBNYV (Figure 5C). The identified site is in agreement with the cognate *ori* of this virus (Figure 5C) [2]. To expand on this further, we used this feature to predict the *ori* sequence of a virus related to phylum *Cressdnaviricota* that has a somewhat more complicated genome structure than current members of the phylum, the prototypical crucivirus Boiling Springs Lake RNA-DNA Hybrid Virus (BSL RDHV) [22]. Because we don’t have any cleavage profiling data for BSL RDHV, we selected the closely related HUH-tag PCV2 to identify cleavage sites across its genome. Even with the increased size and complexity of the BSL RDHV genome, our tool’s prediction is in strong agreement with the cognate *ori* of this virus as well (Figure 5D). Overall, we find these features to have niche utility in HUH-tag biotechnology applications and in HUH-endonuclease basic science and we have implemented them into HUHgle for broad use.

**Figure 5.**
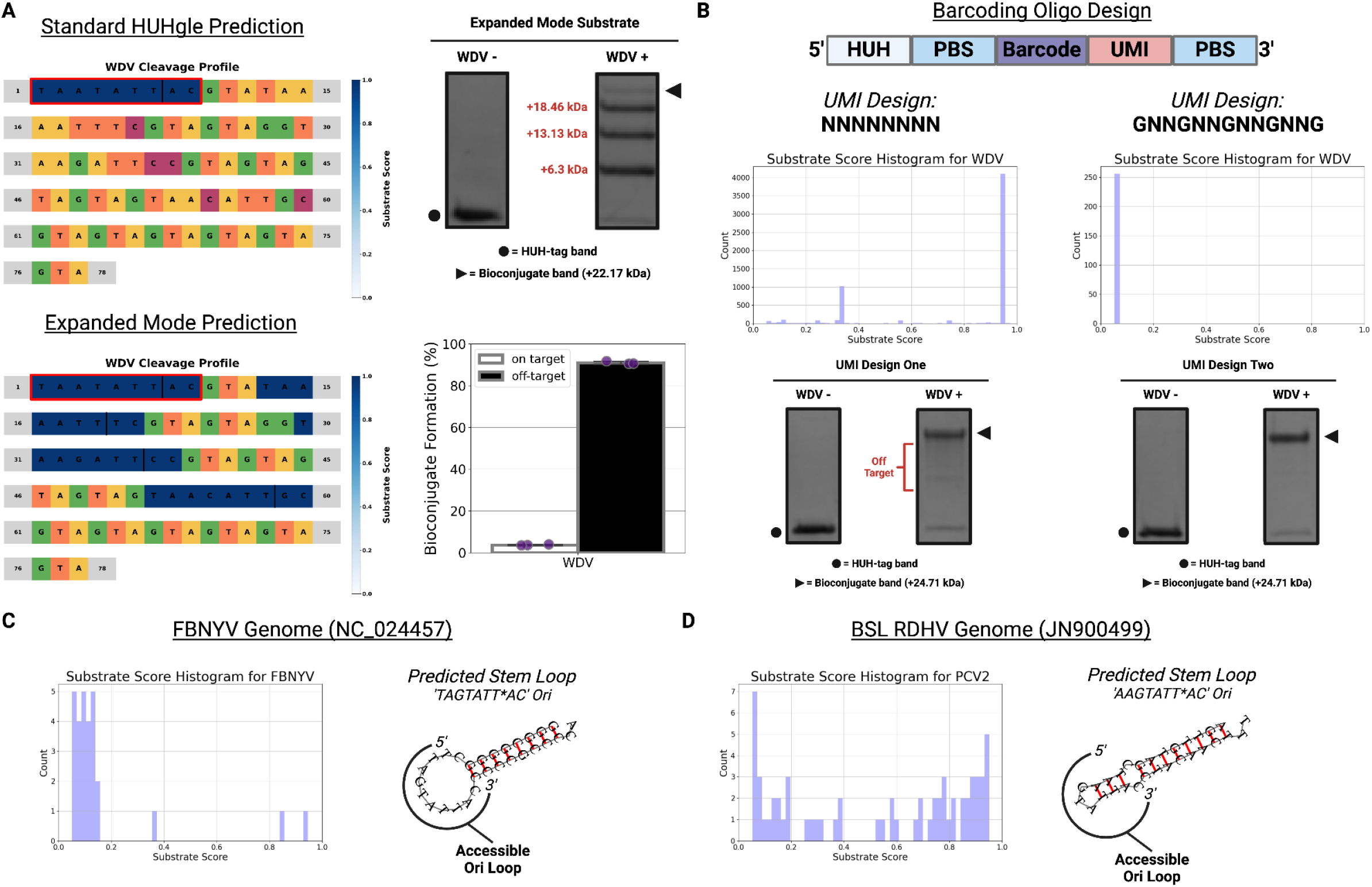
HUHgle bonus features. Subfigure (A) shows comparative HUHgle substrate interaction plots using both standard and expanded mode on a substrate designed to contain a number of cryptic cleavage sites (left), *in vitro* HUH-tag bioconjugation reactions with this cryptic substrate (top right), and a barplot quantifying this reaction (bottom right). Subfigure (B) shows a schematic of the designed barcoding oligonucleotide with an HUH cleavage site (HUH), two primer binding sites (PBS) flanking the barcode and Universal Molecular Identifier (UMI) (top), the output histograms of the bonus feature that enabled HUHgle to interact with ambiguous nucleotide-containing substrates for substrates containing the indicated UMI, and *in vitro* HUH-tag bioconjugation reactions with these substrates visualized via SDS-PAGE. The histograms indicate the number of cleavage sites binned by substrate score for the sequences that the ambiguous substrate codes for. Subfigures (C) and (D) show the output from the HUHgle bonus feature that enables interaction with a .FASTA file and identification of the origin of replication from viruses in phylum *Cressdnaviricota* from the genomes of FBNYV and BSL RDHV, respectively. The histograms indicate the number of cleavage sites binned by substrate score in the entered file and the stemloops images are generated by HUHgle using ViennaRNA on the identified origin of replication-containing stemloop from the indicated virus.

### Conclusions

HUH-tags are a straightforward, site-specific, cost-effective, and chemistry-free way to covalently link proteins of interest to ssDNA. To further enhance the accessibility of HUH-tags, we have implemented HUHgle as a Google Colab notebook with a simple point-and-click user interface, enabling broad usage of our tool regardless of coding experience. Moreover, HUHgle is designed to facilitate incorporation of additional data as groups continue to perform sequence profiling on HUH-endonucleases, which will serve to further expand their utility in both simple and multiplexed applications. Moreover, recent efforts towards virus discovery have massively expanded our understanding of the potential diversity of phylum *Cressdnaviricota* and our knowledge will likely improve further as these efforts continue and the cost of sequencing decreases [23]. Overall, we suspect that HUH-tags will continue to be developed as functional components of a diversity of synthetic biology and biotechnology applications and HUHgle is well suited to enhance and simplify our ability to engineer high-fidelity enzyme-mediated protein-ssDNA bioconjugation.

## Methods

### HUHgle Plotting

Please see the supporting information for a detailed description of the core and bonus HUHgle functions.

### Molecular cloning

The nuclease domain of our panel HUH-endonucleases were synthesized by Integrated DNA Technologies (IDT) as codon-optimized gene fragments. Plasmids were constructed by assembling HUH-tag constructs into the pTD68 expression vector using the In-Fusion HD Cloning Kit (Takara) and verified with Sanger sequencing (Genewiz).

### Protein expression and purification

Constructs were expressed in BL21(DE3) *E. coli* cells (New England BioLabs). Briefly, cultures transformed with 6xHis-SUMO-HUH-tag plasmid were grown in 1 L of LB broth with 100 ug/mL ampicillin (Genesee Scientific). Cultures grew at 37°C shaking at 220 RPM until they reached an OD_600_ of 0.8 when expression was induced via adding 500 μL of 1 M IPTG (isopropyl-D-1-thio-galactopyranoside, Sigma Aldrich) and incubated shaking for 16 hours at 18°C. Cultures were then centrifuged and resuspended in lysis buffer (50 mM Tris pH 7.5, 300 mM NaCl, 1 mM EDTA) and sonicated for three one minute rounds. Lysed cells were clarified by centrifugation and supernatants were batch bound for one hour with HisPure Ni-NTA agarose beads (ThermoFisher) rocking at 4°C. Samples were purified using gravity flow columns, beads were washed using wash buffer (lysis buffer with 30 mM imidazole), and samples were eluted with elution buffer (lysis buffer with 300 mM imidazole). Following this, elutions were further purified and buffer exchanged using the Superdex 200 increase 10/300 (GE Healthcare) size exclusion chromatography column into lysis buffer.

### In vitro HUH bioconjugation assay

HUH-tag cleavage/covalent linkage of ssDNA substrates (IDT) was performed in final concentrations of 3 μM SUMO-HUH-tag and 3 μM oligo in 50 mM HEPES pH 8.0, 50 mM NaCl, 1 mM DTT, and 1 mM MnCl_2_ for one hour at 37°C unless otherwise noted. Samples were then denatured and run on a 4-20% SDS-PAGE gel (Bio-Rad), stained, destained, and imaged. ImageJ [24] was used to quantify percent bioconjugate formation, which was calculated as follows: ((reacted band intensity)/(non-reacted band intensity + reacted band intensity)) x 100.

## Data/Code availability

All data used to inform HUHgle was generated in a previous study [2] and can be found on our lab website (https://www.mechanosome.org/resources). In order to facilitate wide accessibility, HUHgle has been implemented as a Google Colab notebook (https://colab.research.google.com/drive/1mwRby7ckcqoXltUlceWWdqgjylime4l1). Additionally, HUHgle source code is freely available and can be accessed on GitHub (https://github.com/AdamTSmiley) under an open source license.

### Acknowledgements

W.R.G. received funding from NIH NIGMS (R35 GM119483), A.T.S. and N.S.B. received salary support from the Chemistry-Biology Interface Training Grant (T32GM132029), and M.J.M.A. and K.J.T. received salary support from the Muscle training grant (T32AR007612). All figures were made using BioRender.

## Author Contributions

A.T.S. developed and debugged HUHgle. A.T.S. and N.S.B. purified HUH-tag proteins. A.T.S., A.J.H, and A.C.D.L. performed *in vitro* reactions. M.J.M.A. helped develop the HUHgle bonus features. K.J.T. advised in data analysis. W.R.G. oversaw data analysis and manuscript writing. A.T.S. and W.R.G. prepared the manuscript. All authors contributed to editing and gave approval of the final version.

