## Supplementary information for "HUHgle: An Interactive Substrate Design Tool for Covalent Protein-ssDNA Labeling Using HUH-tags"

### Extended Methods

*Introduction:*
HUHgle is a python-scripted DNA sequence searching and plotting tool designed to make HUH-tag mediated protein-ssDNA applications more intuitive and straightforward for all. In the following section, we will describe some of the core components of the HUHgle script at a high level. Please see the HUHgle Colab notebook (https://colab.research.google.com/drive/1mwRby7ckcqoXltUlceWWdqgjylime4l1) for the commented code.

*Initial Sequence Filtering:*
Upon entering a DNA sequence of interest, HUHgle searches the string and filters out sequences that contain non-'ATGC' characters, sequences that are either too long or too short (HUHgle can accommodate sequences ranging from 9 to 300 nt – sequences shorter than 9 nt are incompatible with HUH-tag reactions and sequences longer than 300 nt are challenging to properly visualize in the interactive plots), and, in the context of entering multiple substrates, sequences that are identical to one or more entered sequence(s).

*Substrate Scoring Metric:*
HUHgle searching and plotting is based on a 'substrate score' metric, which indicates how efficiently the substrate will be acted upon by a selected HUH-tag. The specificity profiling data used in HUHgle was generated in a previous study using a novel deep sequencing assay called HUH-seq [1]. Simply, our substrate score is the percent reduction value of a given k-mer when comparing a reacted library to a reference library calculated as (reference count - treatment count/ reference count). Because the reaction conditions of HUH-seq were substoichiometric (300 nM substrate library: 3 µM HUH-tag), the relationship between a given k-mer's percent reduction value and total bioconjugate yield is weakly predictive in the case of some viral families (specifically nanovirus, see Figure 1C) but always correlative, we opted to change the nomenclature to substrate score to enhance clarity. Importantly, HUHgle only considers sequences with substrate scores of 0.05 or higher to be authentic – anything lower is automatically assessed as a score of 0.

*Single HUH-tag Recommendation:*
After entering an appropriate DNA substrate sequence, users are prompted to select an HUH-tag from the provided panel of eight nucleases, or HUHgle can analyze the substrate and recommend its top three choices based on minimizing predicted interaction with the queried sequence in

order to limit off-target product formation. Briefly, HUHgle makes its recommendation by evaluating the queried sequence against the cleavage database for each of the panel of eight HUH-tags. HUHgle exhaustively searches the substrate for sites of interaction and calculates cumulative substrate score values for each nuclease by adding the scores of each identified site of interaction together for a given substrate/HUH-tag combination. Following this, HUHgle recommends its top three HUH-tags based on minimizing the cumulative substrate score. If no sites of interaction are found in the queried sequence for any member of the panel of HUH-tags HUHgle will report that no cleavage is detected and it will return the entire panel of HUH-tags to select from. Furthermore, if cumulative substrate scores are within 5% of each other throughout all members of the HUH-tag panel HUHgle will report that cleavage is comparable across all available HUH-tags and it will return the entire panel of HUH-tags to select from.

*Orthogonality Scoring and Recommendation of HUH-tag Combinations:*
In the context of two- or three-way orthogonal labeling, we have established a robust framework for evaluating and recommending HUH-tag combinations (pairs or triplets) based on their interaction with specific sequences. This framework is based on an 'orthogonality score' metric, designed to reward nuclease combinations with a large orthogonal sequence pool and punish combinations with unevenness (i.e., orthogonal combinations where the majority of mutually exclusive sequences come from one of the members). Importantly, a given substrate is only considered an orthogonal sequence if it has a score of 0.5 (upper threshold) or greater for one HUH-tag and a score of 0.1 (lower threshold) or lower for the other HUH-tag(s) that make up the combination. Orthogonality score is determined by first identifying these orthogonal sequences (as defined by the thresholds) and then calculating as follows:

*Orthogonality Score = C - D*

Variable *C* represents the product of the high-scoring sequence counts, which is calculated by multiplying the number of sequences scored above the upper threshold for the orthogonal sequences across each HUH-tag in the combination being evaluated. Variable *D* represents the sum of the high-scoring sequence counts, which is calculated by adding the number of sequences scored above the upper threshold for the orthogonal sequences across each HUH-tag in the combination being evaluated.

Variable *C* rewards large numbers of high-scoring sequences across the combination of HUH-tags regardless of how evenly dispersed the counts are (i.e., an orthogonal HUH-tag pair with 10 high-scored orthogonal sequences each has the same value as an orthogonal HUH-tag pair in which one nuclease has 100 high-scored orthogonal sequences and the other has only 1). The subtraction of variable *D* from variable *C* punishes highly unbalanced combinations where one of the HUH-tags in the combination has the majority of high-scored orthogonal sequences.

HUH-tag recommendations are made by identifying all potential sites of interaction within a queried substrate and calculating a cumulative substrate score for each HUH-tag (or, in the case of two- or three-way labeling schemes, each HUH-tag combination). The three HUH-tags with the lowest cumulative score are identified as the three nucleases most suitable for the substrate of interest in an effort to minimize off-target interaction. For two- or three-way labeling, this same cumulative score system is used in conjunction with orthogonality score to identify the highest scoring combination while minimizing potential off-target interaction.

*Interactive HUHgle Plotting:*
Once the substrate sequence(s) and HUH-tag(s) are defined, HUHgle proceeds with its core function of generating an interactive plot that indicates the location and efficiency of predicted sites of interaction for the selected HUH-tag on the substrate of interest. The plotting function makes use of the matplotlib library and custom logic to plot and color-code the sequence. A-, T-, G-, and C-bases are all colored uniquely, and identified sites of interaction are recolored to a monochromatic gradient indicative of substrate score as defined by a legend to the right of the plot. Importantly, HUHgle plots are interactive and allow users to click directly on a base of interest to cycle through the other options. Each alteration is automatically reassessed and replotted, enabling users to engineer high fidelity ssDNA substrates rapidly and efficiently through a quick click-based modification of the underlying sequence. When more than one substrate or HUH-tag is selected, buttons populate above the HUHgle plot that enable the user to cycle through their list of substrates and HUH-tags. Finally, interactive plots are occasionally unstable in the Google Colab environment, and, as such, we have added a 'reset' button that allows users to replot their sequence if issues arise.

*Finalizing Interaction and HUHgle Output:*
HUHgle has a 'Download Plot & Info' button for users to press when finished visualizing and analyzing their substrate. Upon clicking, HUHgle analyzes the finalized substrate (if the user made any modifications using the interactive plot) and returns all identified cleavage motifs, their scores, and their locations. Furthermore, HUHgle uses these sites to calculate the identity and molecular weight of all potential protein-DNA bioconjugation products. The computational tool then prints this information directly to the colab notebook and automatically starts the download of a .ZIP file containing the finalized HUHgle plot as a .PNG file, and all of the information listed above as a .TXT file.

*Bonus Feature One - Expanded Mode:*
Previous works have indicated that while the identity of the -4 and +1 positions in a HUH-tag substrate can be highly variable, they must be able to form a proper Watson-Crick (WC) base pairing for cleavage to occur. When expanded mode is activated, HUHgle extends its predictive capabilities by estimating cleavage efficiencies for substrates containing non-cognate WC base pairings between the -4 and +1 positions. The HUH-seq data that HUHgle is built on was collected using a constant 'AC' dinucleotide in the +1 and +2 positions, limiting our resolution on the WC requirement until recently. When activated, HUHgle iterates through each substrate in the specificity profiling and checks if the base at index 3 (corresponding to the -4 position) is a "T". If so, it creates three new sequences by replacing the bases at indices 3 and 7 (corresponding to the +1 position) with the following combinations: "A" and "T", "G" and "C", and "C" and "G". These new sequences are then assigned the same score as the parent sequence and added to the dictionary.

*Bonus Feature Two - Evaluating Semi-Ambiguous Substrates:*
HUHgle includes a bonus feature that allows users to identify potential sites of interaction on substrates containing ambiguous nucleotides. In order to maintain compatibility with the Colab

environment, HUHgle limits the number of ambiguous nucleotides allowed in a substrate sequence to eight. Upon entry of an ambiguous nucleotide-containing sequence, HUHgle generates a list of all possible substrates and then analyzes each generated sequence for potential cleavage sites based on the selected HUH-tag. Due to the potentially high volume of sequences being analyzed in this method (as many as 65, 536 unique sequences) HUHgle plots its analysis as a histogram of identified sites of interaction binned by substrate score. Upon concluding analyses, HUHgle automatically downloads a .ZIP file containing the histogram as a .PNG file and an output .TXT file that has both summarized and detailed information about the identified motifs and their corresponding substrate score.

*Bonus Feature Three - HUHgle.FASTA:*
HUHgle includes a bonus feature that allows users to identify potential sites of interaction on substrates that are too long to be analyzed as conventional HUHgle plots. In this feature, users upload a .FASTA file of their choosing and HUHgle then pulls out pertinent labeling information and analyzes the sequence for potential sites of interaction. It returns the identity, location, and substrate score of each of the sites of interaction that it identifies and plots them as a histogram of sequences binned by substrate score. Importantly, this bonus feature is incompatible with substrates containing ambiguous nucleotides. Moreover, this bonus feature has additional dropdown menus in its interface that enable users to indicate when their file is meant to be circular and when their file represents a genome from pyhlum *Cressdnaviricota*. When users indicate that their sequence is meant to be circular, HUHgle appends the last 30 bases from the end of the file to the beginning in order to capture any sequences that span the function of the end to the beginning. Furthermore, when users indicate that their .FASTA is from phylum *Cressdnaviricota*, HUHgle is able to make a prediction as to what the cognate origin of replication is based on both identified interaction sites for the selected HUH-tag and on how well the substrate folds into a stemloop – a common characteristic of these sequences from this phylum.

### **Supplemental Tables, Data, & Figures**

*Supplementary Table 1 - Panel of HUH-tags in HUHgle*

| HUH-tag | Viral Species | Viral Family | Accession | Cognate *Ori* |
|---|---|---|---|---|
| PCV2 | Porcine circovirus 2 | Circoviridae | NC_005148 | AAGTATT*AC |
| DCV | Muscovy duck circovirus | Circoviridae | KR491947 | TATTATT*AC |
| FBNYV | Faba bean necrotic yellows virus | Nanoviridae | NC_024457 | TAGTATT*AC |
| BBTV | Banana bunchy top virus | Nanoviridae | KM607712 | TATTATT*AC |
| WDV | Wheat dwarf virus | Geminiviridae | AJ311031 | TAATATT*AC |
| TYLCV | Tomato yellow leaf curl virus | Geminiviridae | AJ489258 | TAATATT*AC |
| CLCV | Cabbage leaf curl virus | Geminiviridae | DQ178612 | TAATATT*AC |
| TGMV | Tomato golden mosaic virus | Geminiviridae | JF694490 | TAATATT*AC |

*Supplementary Table 2 - Substrates Analyzed in the Manuscript*

| Name | Sequence | Purpose | Figures |
|---|---|---|---|
| HUHgle1 | GCTCGTCCGTGGTAATATTACCAATAGGATAATTG | High Activity Control | 1C and S2 |
| HUHgle2 | GCTCGTCCGTGGGAATACTACCAATAGGATAATTG | Medium Activity Control (Max) | 1C and S2 |
| HUHgle3 | GCTCGTCCGTGGCCGTAATACCAATAGGATAATTG | Medium Activity Control (Average) | 1C and S2 |
| HUHgle4 | GCTCGTCCGTGGCCTGTAAACCAATAGGATAATTG | No Activity Control | 1C and S2 |
| HUHgle5 | GCTCGTCCGATAATATTACCATAGGATACATTATACCGCTAATTGTTTTTACCAGCC | Single Labeling | 2 |
| HUHgle5Alt | GCTCGTCCGATAATATAACCATAGGATACATTATACCGCTAATTGTTTTTACCAGCC | Single Labeling Alternative | S3 |
| HUHgle6_1 | GAATTTCTACCGTCGGTCCTT | Double Labeling 1 (DCV) | 3A&C |
| HUHgle6_2 | GTAATATCACGGTATT | Double Labeling 1 (TGMV) | 3A&C |
| HUHgle7 | GCTCGTCCGGCATTTATACCGTCGGTCCTTTAAAACTACCGGTATT | Double Labeling 2 (BBTV and CLCV) | 3D&G |
| HUHgle8_1 | GCTCGTCCGGCAGAATTACCGTCGGTCCTTGGA | Triple Labeling 1 (BBTV) | 4A&B |
| HUHgle8_2 | GCTCGTCCGTCATAATACCGGTATT | Triple Labeling 1 (DCV) | 4A&B |
| HUHgle8_3 | GCTCGTCCGGTGTAATATCACCGTT | Triple Labeling 1 | 4A&B |

| | | | |
|---|---|---|---|
| | | (WDV) | |
| HUHgle9 | TCGTCCGGCAGAATTACCGTCGGTCCTTGGATGATAATACCGGTATTTAATATCACCGTT | Triple Labeling (BBTV, DCV, TGMV) | S4 |
| HUHgle10 | TAATATTACGTATAAAATTTCGTAGTAGGTAAGATTCCGTAGTAGTAGTAGTAACATTGCGTAGTAGTAGTAGTAGTA | Expanded Mode Example | 5A |
| HUHgle11 | TTATAATATTACCCCGCGTGATCCAGATACATCGTAGTTGCACGAAATTTTNNNNNNNNNTTAAGTGCGTAGATCAAGGACAGGTTCGTTT | Standard Barcoding Example | 5B |
| HUHgle12 | TTATAATATTACCTGATCCAGATACATCGTAGTTGCACGAAATTTTGNNGNNGNNGNNGTTAAGTGCGTAGATCAAGGACAGGTTCGTTT | High Fidelity Barcoding Example | 5B |

*Supplementary Data 1 - Amino Acid Sequences for Panel of HUH-tags*

>PCV2
PSKKNGRSGPQPHKRWVFTLNNPSEDERKKIRDLPISLFDYFIVGEEGNEEGRTPHLQGFANFVKKQTFNKVKWYLGARCHIEKAKGTDQQNKEYCSKEGNLLMECGAPRSQGQR

>DCV
MAKSGNYSYKRWVFTINNPTFEDYVHVLEFCTLDNCKFAIVGEEKGANGTPHLQGFLNLRSNARAAALEESLGGRAWLSRARGSDEDNEEYCAKESTYLRVGEPVSKGRSS

>FBNYV
MARQVICWCFTLNNPLSPLSLHDSMKYLVYQTEQGEAGNIHFQGYIEMKKRTSLAGMKKLIPGAHFEKRRGTQGEARAYSMKEDTRLEGPWEYGEFVP

>BBTV
MARYVVCWMFTINNPTTLPVMRDEIKYMVYQVERGQEGTRHVQGYVEMKRRSSLKQMRVFFPGAHLEKRKGSQEEARSYCMKEDTRIEGPFEFG

>WDV
MASSSTPRFRVYSKYLFLTYPQCTLEPQYALDSLRTLLNKYEPLYIAAVRELHEDGSPHLHVLVQNKLRASITNPNALNLRMDTSPFSIFHPNIQAAKDCNQVRDYITKEVDSDVNTAEWGTFVAVSTPGRKDRDAD

>TYLCV
MPRLFKIYAKNYFLTYPNCSLSKEEALSQLKKLETPTNKKYIKVCKELHENGEPHLHVLIQFEGKYQCKNQRFFDLVSPNRSAHFHPNIQAAKSSTDVKTYVEKDGNFIDFGVSQIDGRSARGGQQSANDAYAEAL

>CLCV

MPRNPKSFRLAARNIFLTYPQCDIPKDEALQMLQTLSWSVVKPTYIRVAREEHSDGFPHLHCLIQL
SGKSNIKDARFFDITHPRRSANFHPNIQAAKDTNAVKNYITKDGDYCESG

>TGMV
MPSHPKRFQINAKNYFLTYPQCSLSKEESLSQLQALNTPINKKFIKICRELHEDGQPHLHVLIQFEG
KYCCQNQRFFDLVSPTRSAHFHPNIQRAKSSSDVKTYIDKDGDTLVWGEFQVDGRSA

*Supplementary Figures Section*

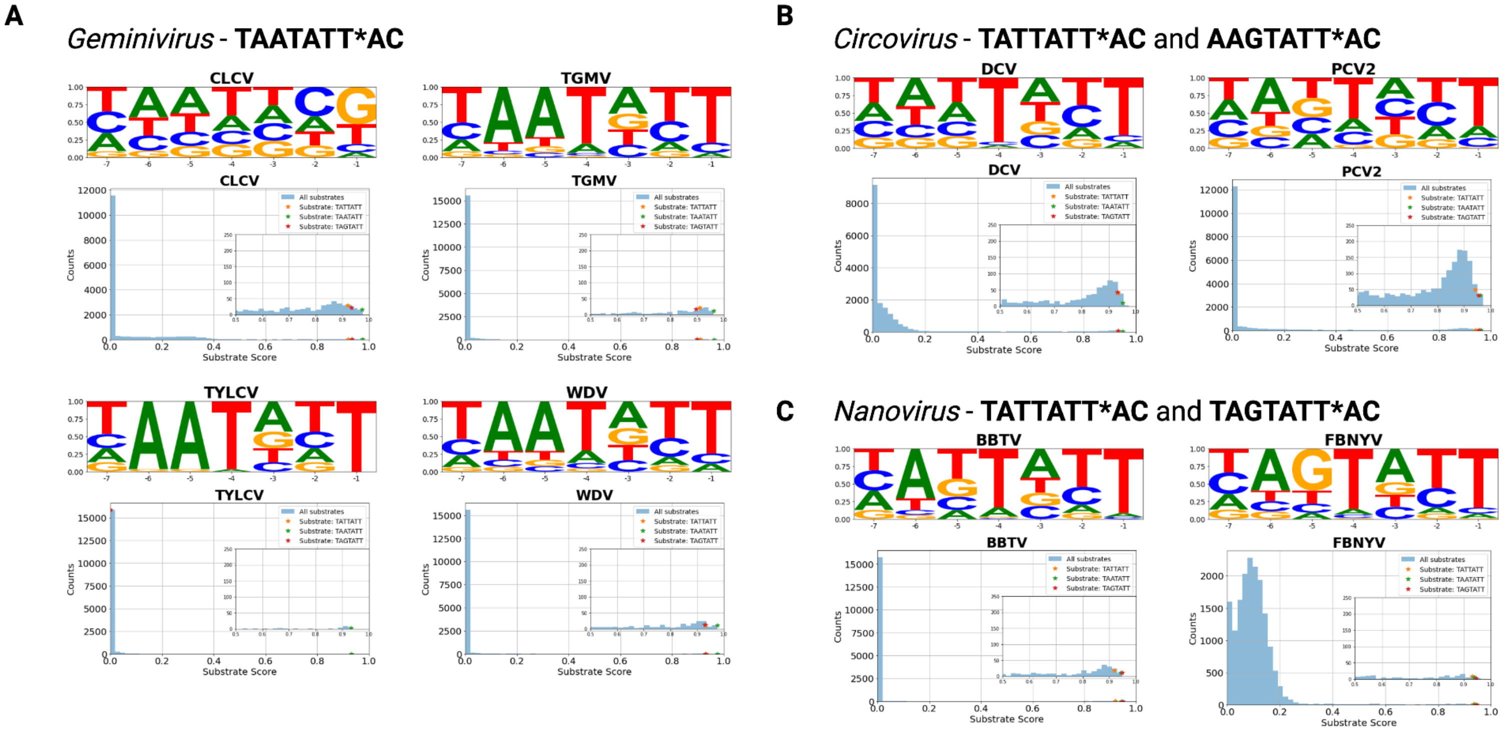


**Figure S1.** *Sequence Logos and Substrate Scores for HUHgle HUH-tag panel*. Subfigures (A), (B), and (C) show sequence specificity logos (top) and histograms of the sequences from specificity profiling binned by substrate score (bottom) separated into pertinent viral families, gemini-, circo-, and nanovirus, respectively. The sequence logos are made using data from the HUH-seq sequence profiling assay and they are indicative of nucleotide preference across the substrate. Only sequences with substrate scores above 0.25 were used to produce these figures. The histograms are made using the entirety of this data, they contain inset graphs for better resolution of sequences scored at 0.5 or above, and substrates corresponding to common endogenous cleavage sites of these enzymes are highlighted with colored stars.

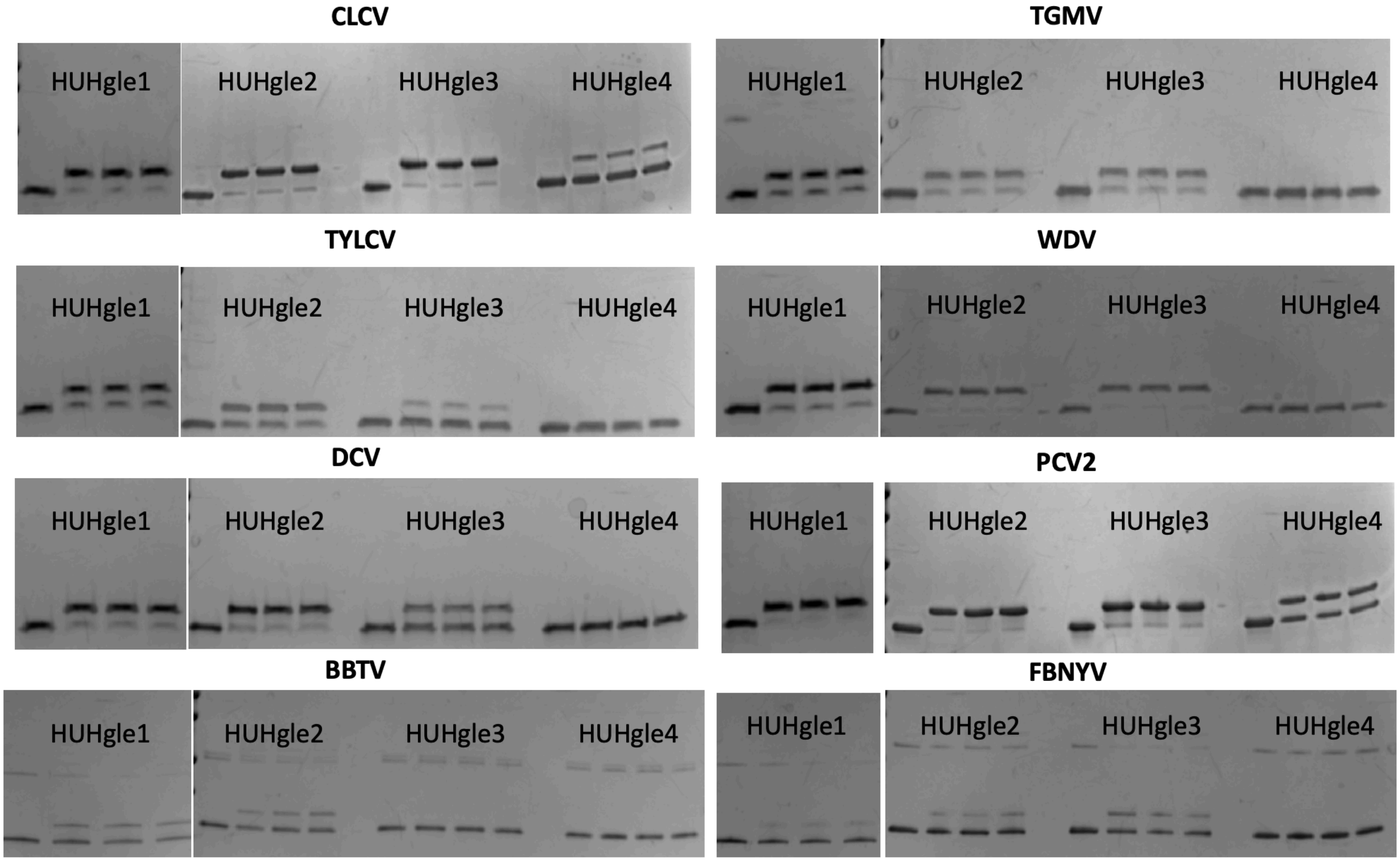


**Figure S2.** *Validation Cleavage Gels.* Gels of *in vitro* HUH-tag bioconjugation reactions analyzed in the scatterplot in figure one visualized via SDS-PAGE. The substrates HUHgle1, HUHgle2, HUHgle3, and HUHgle4 (see supplemental table 2) are reacted with the indicated HUH-tag in triplicate with a single no reaction control.

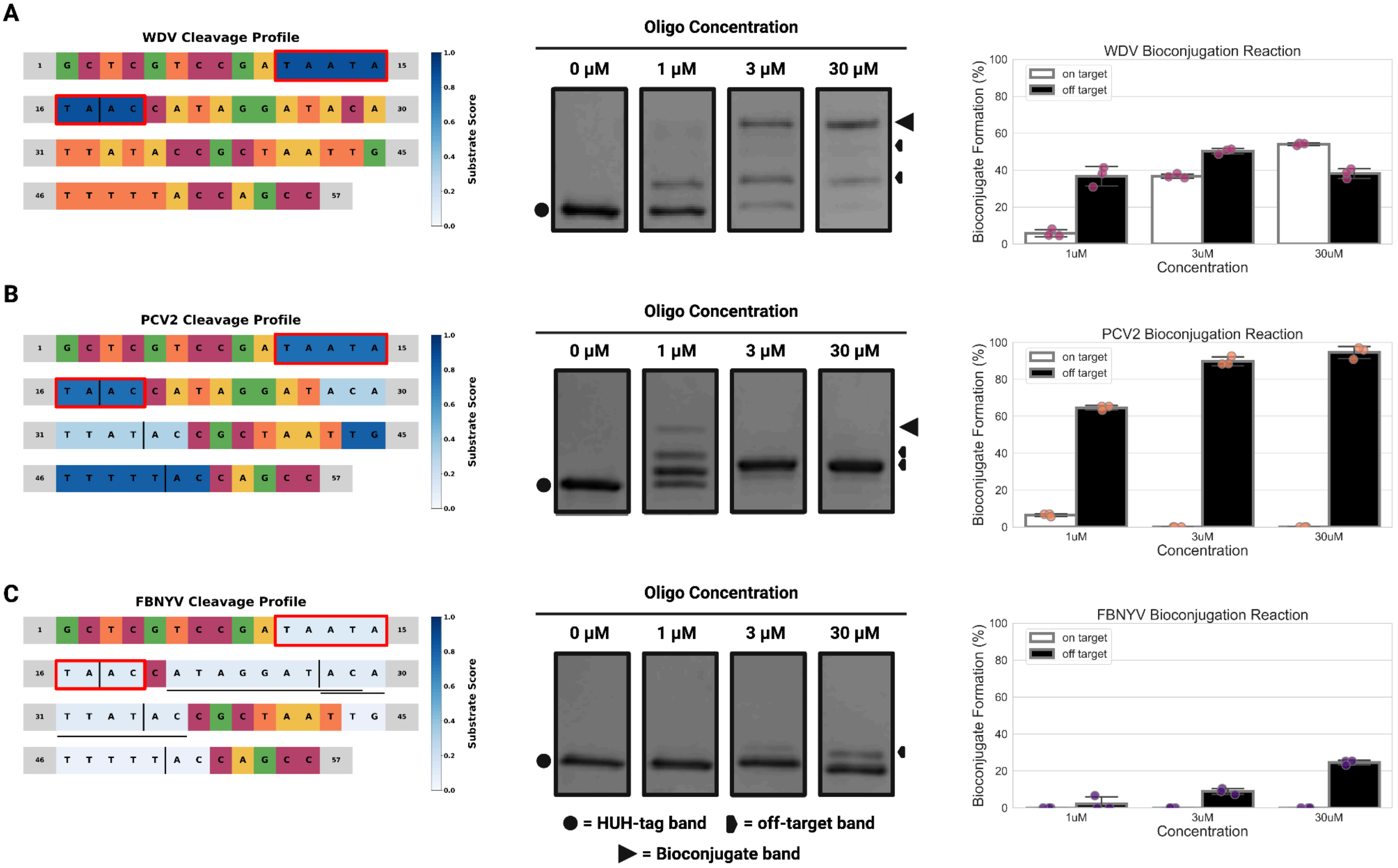


**Figure S3.** *HUHgle predictions in one-way bioconjugation with a sub-optimal designed site:* Subfigures (A), (B), and (C) show HUHgle substrate interaction plots (left), *in vitro* HUH-tag bioconjugation reactions visualized via SDS-PAGE (center), and barplots quantifying these reactions (right) for the HUH-tags WDV (A), PCV2 (B), and FBNYV (C) on the substrate *HUHgle5Alt* (see supporting information). Reactions were performed in final concentrations of 3 µM HUH-tag and indicated concentration of oligo in 50 mM HEPES pH 8.0, 50 mM NaCl, 1 mM DTT, and 1 mM $MnCl_2$ for one hour at 37°C. HUH-tag bands, on-target bioconjugation bands, and the most prominent off-target bioconjugation are indicated with unique black characters – see the legend at the bottom of the figure.

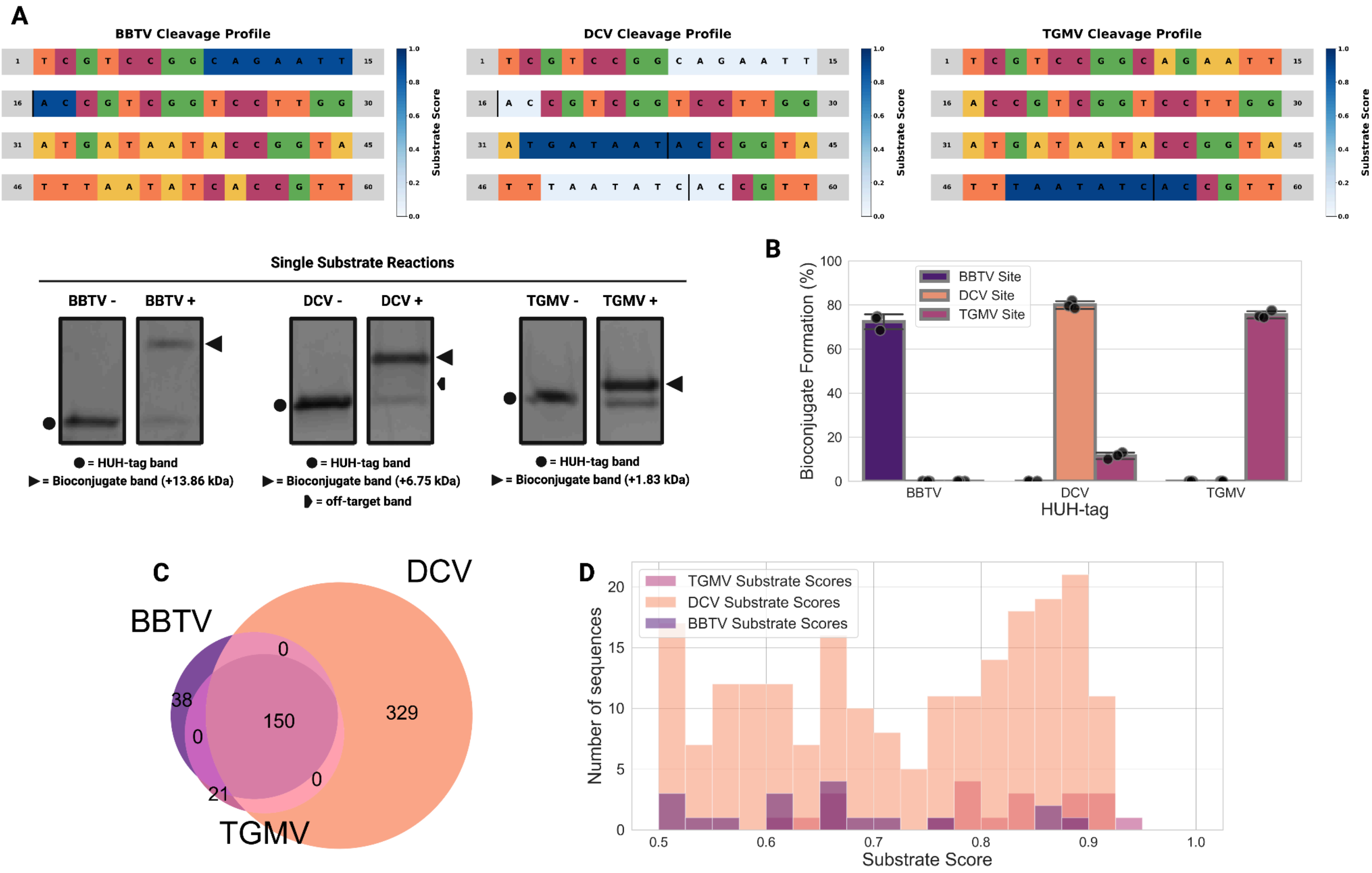


**Figure S4.** *HUHgle predictions in three-way orthogonal bioconjugation.* Subfigure (A) shows HUHgle substrate interaction plots indicating cleavage on a designed substrate containing three orthogonal cleavage sites (top), and *in vitro* HUH-tag bioconjugation reactions with this substrates visualized via SDS-PAGE (bottom) for the orthogonal combination of the HUH-tags BBTV, DCV, and TGMV. Subfigure (B) shows a barplot quantifying these reactions for the labeled HUH-tag and orthogonal substrate cleavage site. Subfigure (C) shows a Venn diagram indicating the number of orthogonal and non-orthogonal substrates scoring above 0.5 across the indicated set of HUH-tags. Subfigure (D) shows a histogram that indicates the number of exclusively orthogonal substrates binned by substrate score across the indicated set of HUH-tags. Reactions were performed in final concentrations of 3 µM HUH-tag and indicated concentration of oligo in 50 mM HEPES pH 8.0, 50 mM NaCl, 1 mM DTT, and 1 mM $MnCl_2$ for 24 hours at 37°C.
